# Abundant secreted hydrolytic enzymes and secondary metabolite gene clusters in genomes of the *Botryosphaeriaceae* reflect their role as important plant pathogens

**DOI:** 10.1101/2021.01.22.427741

**Authors:** JH Nagel, MJ Wingfield, B Slippers

## Abstract

The *Botryosphaeriaceae* are important plant pathogens, but unique in their ability to establish asymptomatic infections that persist for extended periods in a latent state. In this study, we used comparative analyses to consider elements that might shed light on the genetic basis of the interactions of these fungi with their plant hosts. For this purpose, we characterised secreted hydrolytic enzymes, secondary metabolite biosynthetic gene clusters and considered general trends in genomic architecture using all available *Botryosphaeriaceae* genomes, and selected Dothideomycetes genomes. The *Botryosphaeriaceae* genomes were rich in carbohydrate-active enzymes (CAZymes), proteases, lipases and secondary metabolic biosynthetic gene clusters (BGCs) compared to other Dothideomycete genomes. The genomes of *Botryosphaeria, Macrophomina, Lasiodiplodia* and *Neofusicoccum*, in particular, had gene expansions of the major constituents of the secretome, notably CAZymes involved in plant cell wall degradation. The *Botryosphaeriaceae* genomes were shown to have moderate to high GC contents and most had low levels of repetitive DNA. The genomes were not compartmentalized based on gene and repeat densities, but genes of secreted enzymes were slightly more abundant in gene-sparse regions. The abundance of secreted hydrolytic enzymes and secondary metabolite BGCs in the genomes of *Botryosphaeria, Macrophomina, Lasiodiplodia*, and *Neofusicoccum* were similar to those in necrotrophic plant pathogens, but also endophytes of woody plants. The results provide a foundation for future comparative genomic analyses and hypothesis to explore the mechanisms underlying *Botryosphaeriaceae* host-plant interactions.

## 1. Introduction

Secreted hydrolytic enzymes and fungal toxins play crucial roles in enabling fungal pathogens to establish successful infections on their plant hosts. Among the secreted proteins, carbohydrate-active enzymes (CAZymes), protease and lipases are important for nutrient acquisition, as well as for the breakdown, manipulation or circumvention of host defences (Carlile *et al*. 2000; Christensen and Kolomiets 2011; De Jonge and Thomma 2009; Haridas *et al*. 2020; Jia *et al*. 2000; Ohm *et al*. 2012; Voigt *et al*. 2005; Zhao *et al*. 2013). Fungal toxins are a diverse group of compounds and those most commonly found in fungal pathogens include polyketides, non-ribosomal peptides, terpenes and indole alkaloids (Keller *et al*. 2005). These toxins are secondary metabolites that induce plant cell death and for this reason, necrotrophic plant pathogens usually possess greater numbers of genes involved in secondary metabolite synthesis than biotrophic pathogens (Howlett 2006).

The genomes of many fungal and Oomycetes plant pathogens, especially those rich in repetitive elements, are not homogenous, but rather compartmentalized into repeat-rich, gene sparse regions and repeat poor, gene dense regions (Dong *et al*. 2015; Frantzeskakis *et al*. 2019; Ma *et al*. 2010; Raffaele *et al*. 2010; Raffaele and Kamoun 2012; Rouxel *et al*. 2011). Genes localized to repeat-rich, gene sparse regions also have a higher rate of mutation and are often under stronger selective pressure (Möller and Stukenbrock 2017; Raffaele and Kamoun 2012; Stukenbrock *et al*. 2010). This has given rise to a phenomenon referred to as ‘two-speed’ genomes, due to the stark differences in evolutionary rates between the two different types of genomic regions.

Fungi residing in the *Botryosphaeriaceae* include important plant pathogens. These fungi mostly cause diseases of woody plant species and they can impact negatively on the health of many economically and ecologically significant plant species (Mehl *et al*. 2013; Slippers and Wingfield 2007). The *Botryosphaeriaceae* infect a wide range of plant hosts, most notably grapevine (Urbez-Torres 2011), pome and stone fruits (Slippers *et al*. 2007), plantation forest trees such as *Eucalyptus* spp., *Pinus* spp. and *Acacia mangium* (Alves *et al*. 2013; Mohali *et al*. 2007; Rodas *et al*. 2009), as well as plants in their native habitats (Jami *et al*. 2014; Jami *et al*. 2017; Marincowitz *et al*. 2008; Pavlic *et al*. 2008). Many of these fungi (e.g. *B. dothidea, M. phaseolina, Lasiodiplodia theobromae, Neofusicoccum parvum*) have wide host ranges, however, some species (e.g. *Diplodia sapinea*) have narrower host ranges or are even very host-specific such as in the case of *Eutiarosporella darliae, E. pseudodarliae* and *E. tritici-australis*. Many species of *Botryopshaeriaceae* are also known to occur endophytically in asymptomatic plant tissues or to have a latent pathogenic phase, where they inhabit their plant hosts in the absence of symptoms and cause disease only after the onset of stress, such as drought, frost or hail damage (Bihon *et al*. 2011; Slippers and Wingfield 2007).

A few recent studies have investigated secreted proteins and secondary metabolites in species of the *Botryosphaeriaceae*. Proteomic studies analyzing the secreted proteins of *Diplodia seriata* (Cobos *et al*. 2010) and *D. corticola* (Fernandes *et al*. 2014) identified secreted proteins involved in pathogenesis. Studies of grapevine pathogens also predicted secreted CAZymes and genes involved in the production of secondary metabolites of *D. seriata* and *Neofusicoccum parvum* (Massonnet *et al*. 2016; Morales-Cruz *et al*. 2015). Secondary metabolite biosynthetic gene clusters (BGCs) have also been shown to play a role in host range determination, e.g. in *E. darliae* and *E. pseudodarliae* causing white grain disorder, where the presence of a secondary metabolite biosynthetic gene cluster allows woody hosts to be infected (Thynne *et al*. 2019). Despite the many publicly available genomes of species of *Botryosphaeriaceae* (Blanco-Ulate *et al*. 2013; Islam *et al*. 2012; Liu *et al*. 2016; Marsberg *et al*. 2016; Massonnet *et al*. 2016; Morales-Cruz *et al*. 2015; Robert-Siegwald *et al*. 2017; van der Nest *et al*. 2014), no comprehensive comparative studies have been undertaken using these genomes; neither have analyses been conducted to characterise secreted proteins and secondary metabolites in most these fungi. Such studies are also hampered by the lack of publically available genome annotations.

The manner in which plants interact with beneficial microorganisms, while at the same time restricting the negative effects of pathogens, is an important and intriguing question in plant biology (Southworth 2012). One proposed model referred to as the ‘balanced antagonism model’ (Schulz and Boyle 2005) holds that endophytism is a result of both the host plant and the fungus employing antagonistic measures against each other, in such a way that neither overwhelms the other. Disruption of this balance either results in the pathogen causing disease or in the host plant successfully killing the fungus. The model thus predicts that known endophytic species should have similar genetic repertoires to their closely related plant pathogenic relatives. This appears to be the case when considering recent comparative genomics studies conducted on endophytic fungi (Hacquard *et al*. 2016; Schlegel *et al*. 2016; Xu *et al*. 2014; Yang *et al*. 2019), although some endophytic species, e.g. *Xylonia heveae* had fewer CAZymes than expected and were more similar to mutualistic species (Gazis *et al*. 2016). Indeed, the above-mentioned endophytes (other than *X. heveae*) commonly had high numbers of plant cell wall degrading enzymes and secondary metabolite genes.

Despite their ubiquity as endophytes and their importance as latent pathogens, very little is known regarding how *Botryosphaeriaceae* species interact with their plant hosts at a molecular level. Key questions in this regard relate to the secreted hydrolytic enzymes and secondary metabolic biosynthesis genes present in their genomes. Based on the results of previous studies on Ascomycetes that are endophytes of woody plants, we have hypothesised that these genes and gene clusters in the *Botryosphaeriaceae* will resemble those of closely related plant pathogens. To test this hypothesis, we compared the secreted hydrolytic enzyme and secondary metabolite genes of *Botryosphaeriaceae* species with those of other Dothideomycetes. We also characterised the genome architecture of the *Botryosphaeriaceae* in terms of gene density, repeat content and prevalence of repeat-induced point mutations (RIP), and considered how these associate with secreted hydrolytic enzymes and secondary metabolite BGCs.

## 2. Materials and Methods

### 2.1. Genomic data

All available, published *Botryosphaeriaceae* genomes were retrieved from public databases (NCBI and JGI). Additionally, we sequenced and assembled 12 genomes representing three *Lasiodiplodia* spp. and five *Neofusicoccum* spp. (Table 1). To standardize protein annotations for downstream application, all of the above *Botryosphaeriaceae* genomes were annotated using the same pipeline described below. Additionally, the genomes and protein annotations of 41 Dothideomycetes and *Aspergillus nidulans* (Eurotiales), which had both genomic sequences and annotated protein sequences available on NCBI, were retrieved (Table 1). These genomes were used for comparative purposes in the phylogenomic analyses, hydrolytic enzyme and secondary metabolite BGC analyses and in the statistical clustering analyses, described below.

**Table 1.** List of genome sequences used in this study*

| Species | Reference collection/Isolate number | Assembly Size (Mbp) | Number of gene models | Genome accession number | Reference |
| --- | --- | --- | --- | --- | --- |
| Dothideomycetes |  |  |  |  |  |
| Botryosphaerales |  |  |  |  |  |
| *Botryosphaeria kuwatsukai* | LW030101 | 47.39 | 11 278 | MDSR01000000 | Lui et al. 2016 |
| *Botryosphaeria dothidea* | CMW8000 | 43.50 | 11 368 | http://genome.jgi.doe.gov/Botdo1_1/Botdo1_1.home.html | Marsberg et al. 2017 |
| *Diplodia corticola* | CBS112549 | 34.99 | 9 376 | MNUE01000001 | Fernandes et al. 2014 |
| *Diplodia sapinea* | CBS117911 | 36.05 | 9 589 | AXCF00000000 | Bihon et al. 2014 |
|  | CBS138184 | 35.24 | 9 386 | JHUM00000000 | Bihon et al. 2014 |
| *Diplodia scrobiculata* | CBS139796 | 34.93 | 9 204 | LAEG00000000 | Wingfield et al. 2015 |
| *Diplodia seriata* | UCDDS831 | 37.12 | 9 759 | MSZU00000000 | Morales-Cruz et al. 2015 |
|  | F98.1 | 37.27 | 9 832 | LAQI00000000 | Robert-Siegwald et al. 2017 |
| *Eutiarosporella darliae* | 2G6 | 27.27 | 7 904 | GFXH01000000 | Thynne et al. 2019 |
| *Eutiarosporella pseudodarliae* | V4B6 | 26.74 | 7 846 | GFXI01000000 | Thynne et al. 2019 |
| *Eutiarosporella tritici-australis* | 153 | 26.59 | 7 783 | GFXG01000000 | Thynne et al. 2019 |
| *Lasiodiplodia gonubiensis* | **CBS115812** | 41.14 | 10 649 | **RHKH00000000** | Present study |
| *Lasiodiplodia pseudotheobromae* | **CBS116459** | 43.01 | 10 964 | **RHKG00000000** | Present study |
| *Lasiodiplodia theobromae* | **CBS164.96** | 42.97 | 10 961 | **RHKF00000000** | Present study |
|  | CSS01 | 43.28 | 11 017 | RHKB00000000 | Yan et al. 2017 |
| *Macrophomina phaseolina* | MS6 | 48.88 | 10799 | AHHD00000000 | Islam et al. 2012 |
| *Neofusicoccum cordaticola* | **CBS123634** | 45.71 | 12822 | **RHKC00000000** | Present study |
|  | **CBS123638** | 43.56 | 12630 | **RHKD00000000** | Present study |
| *Neofusicoccum kwambonambiense* | **CBS123639** | 44.17 | 12839 | **RHKE00000000** | Present study |
|  | **CBS123642** | 44.21 | 12904 | **RKSS00000000** | Present study |
| *Neofusicoccum parvum* | **CMW9080** | 41.41 | 12870 | **RHJX00000000** | Present study |
|  | **CBS123649** | 42.16 | 12453 | **RHJY00000000** | Present study |
|  | UCRNP2 | 42.52 | 12691 | AORE00000000 | Blanco-Ulate et al. 2013 |
| *Neofusicoccum ribis* | **CBS115475** | 43.18 | 12708 | **RHJZ00000000** | Present study |
|  | **CBS121.26** | 43.12 | 12733 | **RHKA00000000** | Present study |
| *Neofusicoccum umdonicola* | **CBS123644** | 42.29 | 12816 | **RHKB00000000** | Present study |
| Capnodiales |  |  |  |  |  |
| *Acidomyces richmondensis* | meta | 26.82 | 10338 | JOOL00000000 | Mosier et al. 2016 |
| *Baudoinia panamericana* | UAMH 10762 | 21.88 | 10508 | AEIF00000000 | Ohm et al. 2012 |
| *Cercospora berteroeae* | CBS538.71 | 33.89 | 11903 | PNEN00000000 | De Jonge et al. 2018 |
| *Cercospora beticola* | 09-40 | 37.06 | 12463 | LKMD00000000 | De Jonge et al. 2018 |
| *Cercospora zeina* | CMW25467 | 40.76 | 10193 | MVDW00000000 | Wingfield et al. 2017 |
| *Dothistroma septosporum* | NZE10 | 30.21 | 12415 | AIEN00000000 | Ohm et al. 2012 |
| *Pseudocercospora eumusae* | CBS 114824 | 47.12 | 12632 | LFZN01000000 | Chang et al. 2016 |
| *Pseudocercospora fijiensis* | CIRAD86 | 29.98 | 13066 | AIHZ00000000 | Ohm et al. 2012 |
| *Pseudocercospora musae* | CBS 116634 | 60.44 | 13129 | LFZO00000000 | Chang et al. 2016 |
| *Ramularia collo-cygni* | URUG2 | 32.25 | 11612 | FJUY00000000 | McGrann et al. 2016 |
| *Sphaerulina musiva* | SO2202 | 29.35 | 10233 | AEFD00000000 | Ohm et al. 2012 |
| *Zymoseptoria tritici* | IPO323 | 39.69 | 10963 | ACPE00000000 | Goodwin et al. 2011 |
| Dothideales |  |  |  |  |  |
| *Aureobasidium namibiae* | CBS 147.97 | 25.43 | 10259 | AYEM00000000 | Gostinčar et al. 2014 |
| *Aureobasidium subglaciale* | EXF-2481 | 25.80 | 10792 | AYYB00000000 | Gostinčar et al. 2014 |
| Hysteriales |  |  |  |  |  |
| *Hysterium pulicare* | CBS 123377 | 38.43 | 12352 | AJFK00000000 | Ohm et al. 2012 |
| *Rhytidhysterium rufulum* | CBS 306.38 | 40.18 | 12117 | AJFL00000000 | Ohm et al. 2012 |
| Myriangiales |  |  |  |  |  |
| *Elsinoe australis* | NL1 | 23.34 | 9223 | NHZQ00000000 | Zhao et al. 2020 |
| Mytilinidiales |  |  |  |  |  |
| *Lepidopterella palustris* | CBS 459.81 | 45.67 | 13861 | LKAR00000000 | Peter et al. 2016 |
| Pleosporales |  |  |  |  |  |
| *Alternaria alternata* | SRC11rK2f | 32.99 | 13466 | LXPP00000000 | Zeiner et al. 2016 |
| *Ascochyta rabiei* | ArDII | 34.66 | 10596 | JYNV00000000 | Verma et al. 2016 |
| *Bipolaris maydis* | ATCC 48331 | 32.93 | 12705 | AIHU00000000 | Ohm et al. 2012 |
| *Bipolaris oryzae* | ATCC 44560 | 31.36 | 12002 | AMCO00000000 | Condon et al. 2013 |
| *Bipolaris sorokiniana* | ATCC 44560 | 34.41 | 12214 | AEIN00000000 | Condon et al. 2013 |
| *Bipolaris victoriae* | FI3 | 32.83 | 12882 | AMCY00000000 | Condon et al. 2013 |
| *Bipolaris zeicola* | 26-R-13 | 31.27 | 12853 | AMCN00000000 | Condon et al. 2013 |
| *Clohesyomyces aquaticus* | CBS 115471 | 49.68 | 15811 | MCFA00000000 | Mondo et al. 2017 |
| *Corynespora cassiicola* | CCP | 44.85 | 17158 | NSJI00000000 | Lopez et al. 2018 |
| *Epicoccum nigrum* | ICMP 19927 | 34.74 | 12025 | NCTX00000000 | Fokin et al. 2017 |
| *Exserohilum turcicum* | Et28A | 43.01 | 11698 | AIHT00000000 | Ohm et al. 2012 |
| *Leptosphaeria maculans* | JN3 | 45.12 | 12469 | FP929064:FP929139 | Rouxel et al. 2011 |
| *Paraphaeosphaeria sporulosa* | AP3s5-JAC2a | 38.46 | 14734 | LXPO00000000 | Zeiner et al. 2016 |
| *Parastagonospora nodorum* | SN15 | 37.21 | 15994 | AAGI00000000 | Hane et al. 2007 |
| *Periconia macrospinoso* | DSE2036 | 54.99 | 18735 | PCYO00000000 | Knapp et al. 2018 |
| *Pyrenophora tritici-repentis* | Pt-1C-BFP | 38.00 | 12169 | AAXI00000000 | Manning et al. 2013 |
| *Stemphylium lycopersici* | CIDEFI 216 | 35.17 | 8997 | LGLR00000000 | Franco et al. 2015 |
| Venturiales |  |  |  |  |  |
| *Verruconis gallopava* | CBS 43764 | 31.78 | 11357 | JYBX00000000 | Teixeira et al. 2017 |
| *incertae sedis* |  |  |  |  |  |
| *Cenococcum geophilum* | 1.58 | 177.56 | 14709 | LKKR00000000 | Peter et al. 2016 |
| *Coniosporium apollinis* | CBS 100218 | 28.65 | 9308 | AJKL00000000 | Teixeira et al. 2017 |
| *Glonium stellatum* | CBS 207.34 | 40.52 | 14277 | LKAO00000000 | Peter et al. 2016 |
| Eurotiomycetes |  |  |  |  |  |
| Eurotiales |  |  |  |  |  |
| *Aspergillus nidulans* | FGSC A4 | 30.28 | 9556 | AACD00000000 | Galagan et al. 2005 |

\*Entries in boldface represent genomes that were sequenced as part of the present study

#### 2.1.1. DNA extraction, genome sequencing and assembly

Cultures of three *Lasiodiplodia* and five *Neofusicoccum* species (Table 1) were inoculated onto cellophane covered 2 % malt extract agar (MEA; Biolab, Merck) and incubated at 22 °C. After five days, mycelium was harvested from the surface of the cellophane using a sterile scalpel and DNA was extracted using a modified phenol/chloroform protocol. Mycelium was ground to a fine powder using liquid Nitrogen and approximately 500 mg of ground mycelium was used for DNA extraction. To the ground mycelia, 18 ml of a 200 mM Tris-HCl (pH 8.0), 150 mM NaCl, 25 mM EDTA (Ethylenediaminetetraacetic acid, pH 8.0) and 0.5% SDS (Sodium dodecyl sulfate) solution and 125 μl of 20 mg/ml Proteinase K was added and incubated at 60 °C for two hours. This was followed by addition of 6 ml of 5M potassium acetate and 30 minutes incubation at 0 °C. Samples were then centrifuged at 5000 g for 20 minutes. The aqueous phase was kept and 24 ml of a 1:1 phenol:chloroform solution was added; samples were then centrifuged as above. Two chloroform washes were performed on the aqueous phase, followed by addition of 100 μl of 10 mg/ml Rnase A and incubated for two hours. DNA was precipitated using one volume isopropanol and centrifuged for 30 minutes. The pellet was cleaned through two 70% ethanol washes and resuspended in 1 x Tris-EDTA buffer.

The extracted DNA was used for paired-end sequencing (average fragment size of 500 bp). All samples were sequenced on an Illumina HiSeq 2500 platform, except for *N. parvum* (isolate CMW9080) that was sequenced on a Miseq platform. The quality of the resulting reads was assessed using FastQC 0.10.1 (Andrews 2010) and low quality and short reads were trimmed or discarded using Trimmomatic 0.30 (Bolger *et al*. 2014). *De novo* genome assembly was performed Velvet 1.2.10 (Zerbino and Birney 2008) and Velvetoptimiser 2.2.5 (Gladman and Seemann 2012). Paired-end reads were used to scaffold the assembly and insert size statistics were determined by Velvet for each genome assembly. Genome assembly summary statistics were calculated with the AssemblyStatsWrapper tool of BBtools 38.00 (Bushnell *et al*. 2017).

#### 2.1.2. Genome annotation

The twelve sequenced genomes described above, as well as the fourteen *Botryosphaeriaceae* genomes retrieved from public databases, were annotated as follows: Custom repeat libraries were constructed for each genome assembly using RepeatModeler 1.0.10 (Smit and Hubley 2017). BRAKER 1.10 (Hoff *et al*. 2015) was used to create trained GeneMark-ET 4.29 (Lomsadze *et al*. 2014) and AUGUSTUS 3.2.3 (Stanke *et al*. 2006) profiles using previously published *N. parvum* transcriptome data (SRR3992643 and SRR3992649) (Massonnet *et al*. 2016). Genomes were annotated through the MAKER2 2.31.8 (Holt and Yandell 2011) pipeline using the custom repeat libraries and the BRAKER trained GeneMark-ET 4.29 and AUGUSTUS profiles. Genomes and genome annotations were assessed for completeness with BUSCO 4.0.5 (Waterhouse *et al*. 2017) using the Ascomycota ortholog library (Creation date 2020-09-10, 1706 core orthologous genes). The annotations for six *Botryosphaeriaceae* genomes availablet on public databases prior to this study were also assessed using BUSCO and compared to those of the annotations generated in this study.

### 2.2. Phylogenomic analyses

To illustrate the relationships between species and genera of the *Botryosphaeriaceae*, as well as the relationship of this family to the rest of the Dothideomycetes, a robust phylogeny was created from the genome data. Single copy core orthologous genes were identified from individual genomes (Table 1) using BUSCO (as described above). The BUSCO genes present in all taxa (207 genes) were selected for further analysis. Each orthogroup was aligned using MAFFT 7.407 (Katoh and Standley 2013) before being concatenated into a single matrix. RAxML 8.2.4 (Stamatakis 2014) was used to perform maximum likelihood phylogenetic inference using the PROTGAMMAAUTO option and a thousand bootstrap replicates were performed. Trees were rooted using sequences from *Aspergillus nidulans*.

### 2.3. Functional annotation

The genome annotation data for the *Botryosphaeriaceae*, the other Dothideomycetes and the outgroup *A. nidulans* were used to perform functional annotation for hydrolytic enzymes and secondary metabolite BGCs. CAZymes were predicted by searching the total predicted proteins of each genome against the CAZy database (Lombard *et al*. 2013) using dbCAN2 (HMMdb release v9.0) (Zhang *et al*. 2018). Only those CAZyme predictions supported by two or more tools (HMMR, DIAMOND, Hotpep) were retained. Proteases and protease inhibitors were predicted by subjecting the predicted protein sequences to a BLASTP (Camacho *et al*. 2009) search against the MEROPS protease database 12.0 (Rawlings *et al*. 2017) using a cut off E-value of 1E-04. Lipases and cutinases were predicted by searching protein sequences against lipase and cutinase hidden Markov model (HMM) profiles retrieved from the Lipase Engineering Database v 3.0 (Fischer and Pleiss 2003) using HMMER 3.1b2 (Eddy 2011). Secondary metabolite BGCs were identified from all annotated genomes. using AntiSMASH v5.1.1 (Blin *et al*. 2019).

The total predicted proteins were also analyzed for the presence of signal peptides, involved in protein secretion. Phobius 1.01 (Käll *et al*. 2007) and SignalP 4.1 (Petersen *et al*. 2011) were used to assess the presence of signal peptides and TMHMM 2.0 (Krogh *et al*. 2001) was used to determine if any transmembrane regions occurred within these proteins. Only proteins with a signal peptide predicted by both Phobius and SignalP, as well as no transmembrane domains outside of the signal peptide predicted by both Phobius and TMHMM were regarded here as predicted secreted proteins.

### 2.4. Analysis of gene family evolution

CAFE (Computational Analysis of gene Family Evolution) v4.2 (Han *et al*. 2013) was used to study the evolution of gene family size in the hydrolytic enzymes and secondary metabolite BGCs. To this end, the phylogeny described above was converted to an ultrametric tree using r8s v1.81 (Sanderson 2003). This was then calibrated by fixing the age of the *Botrosphaeriaceae* to 61 million years (Phillips *et al*. 2019) and constraining the age of the Dothideomycetes to 303-357 million years (Hyde *et al*. 2017). Gene family sizes for the total predicted CAZymes, proteases, lipases and secondary metabolite BGCs (Supplementary File 1) were used as input for the CAFE analyses. CAFE was run using separate lambda (birth) and mu (death) rate parameters. Additionally, two separate rate classes were allowed in that the rate parameters were calculated independently for the *Botryosphaeriaceae* and the remaining Dothideomycete taxa. CAFE was run using a P-value cutoff of 0.01 and Viterbi P-values were calculated to significant expansions/contractions across branches.

### 2.5. Hierarchical clustering and pricipal component analysis

Hierarchical clustering was done to determine the similarity between taxa based on the secreted hydrolytic enzyme classes and secondary metabolite BGC types. The heatmap.2 function of the gplots (Warnes *et al*. 2016) R package was used to perform the analysis. Aditionally, principal component analysis (PCA) was performed using the same functional annotations as used in the hierarchical clustering, with the exception that for CAZymes and proteases the number of genes associated with each family (e.g AA1, A01) was used instead of those of each class (e.g AA, Aspartic). The FactoMineR (Lê *et al*. 2008) R package was used to perform the analysis and Factoshiny (Vaissie *et al*. 2015) was used to generate PCA plots.

### 2.6. Genome architecture

Gene densities were analyzed for each *Botryosphaeriaceae* genome to assess the level of genome compartmentalization. Intergenic distances were used as a measure of gene density, by considering the 5’ and 3’ flanking intergenic regions (FIRs) of each gene. FIR lengths were used for two-dimensional data binning to construct gene density heat maps (Saunders *et al*. 2014) using R (Team 2013). Possible differences in the gene density distributions between the total and secreted CAZymes, proteases and lipases were also considered. We compared the relative amounts (i.e. the percentage) of these genes located in gene-sparse regions. In this case, gene sparse regions were defined as genes with both 5’ and 3’ FIRs larger than 1500 bp, as previously used by Raffaele and Kamoun (2012)

The repeat contents of the *Botryosphaeriaceae* genomes were determined using RepeatMasker (Smit *et al*. 2017) with custom repeat libraries created by RepeatModeler. The presence of TA rich regions and the genes present therein were determined using OcculterCut (Testa *et al*. 2016). Specifically, the numbers of the total secreted genes, the secreted CAZymes, proteases and lipases, as well as the number of secondary metabolite BGCs associated with TA rich regions were determined. The occurrence of RIP, including large regions affected by RIP (LRARs), was determined in each genome using TheRIPper (Van Wyk *et al*. 2019)

## 3. Results

### 3.1 Genome sequencing, assembly and annotation

Nine genomes of *Neofusicoccum* and three genomes of *Lasiodiplodia* species were sequenced using Illumina sequencing. These included two isolates each of *N. cordaticola, N. kwambonambiense, N. parvum* and *N. ribis* were sequenced. A single isolate was sequenced for *L. gonubiensis, L. pseudotheobromae, L. theobromae* and *N. umdonicola*.

*De novo* genome assembly resulted in genome lengths of approximately 43 MB for both *Lasiodiplodia* spp. and *Neofusicoccum* spp. (Table 2). The number of scaffolds/contigs was very variable between the sequenced genomes, but the three *Lasiodiplodia* genomes had a lower number of scaffolds than the *Neofusicoccum* genomes. The *N. parvum* CMW9080 genome that was sequenced on the Miseq platform had a higher degree of fragmentation, as seen from the high total number of scaffolds and a large number of short contigs and scaffolds. The percentage of repetitive elements of each genome was higher on average in the *Neofusicoccum* genomes (6.84%) than in the *Lasiodiplodia* genomes (3.33%).

**Table 2:** Genome statistics of new draft *Botryosphaeriaceae* genomes

|  | #<br>scaffolds | #<br>contigs | Scaffold<br>length<br>(Mb) | Contig<br>length<br>(Mb) | Genome scaffold<br>N50/L50 (#/kb) | Genome contig<br>N50/L50 (#/kb) | Maximum<br>scaffold length<br>(Mb) | Maximum<br>contig length<br>(kb) | % main genome<br>in scaffolds ><br>50 KB | % GC | %<br>repetitive<br>sequences |
| --- | --- | --- | --- | --- | --- | --- | --- | --- | --- | --- | --- |
| *Lasiodiplodia gonubiensis*<br>CBS115812 | 376 | 1578 | 41.14 | 40.97 | 50/234.83 | 267/45.4 | 1.06 | 226.76 | 0.92 | 54.94 | 3.23 |
| *Lasiodiplodia pseudotheobromae*<br>CBS116459 | 403 | 1285 | 43.01 | 42.89 | 48/236.23 | 218/61.88 | 1.03 | 416.59 | 0.91 | 54.82 | 3.26 |
| *Lasiodiplodia theobromae*<br>CBS164.96 | 424 | 1093 | 42.97 | 42.88 | 48/223.84 | 163/81.56 | 1.69 | 591.21 | 0.91 | 54.86 | 3.49 |
| *Neofusicoccum cordaticola*<br>CBS123634 | 1912 | 8698 | 45.71 | 45.10 | 276/47.38 | 1218/10.76 | 680.02 | 143.71 | 0.48 | 55.77 | 7.86 |
| *Neofusicoccum cordaticola*<br>CBS123638 | 2393 | 3252 | 43.56 | 43.44 | 254/51.16 | 386/33.04 | 274.15 | 152.21 | 0.51 | 56.15 | 8.30 |
| *Neofusicoccum kwambonambiense*<br>CBS123639 | 1560 | 2839 | 44.17 | 43.99 | 219/59.44 | 363/34.21 | 344.53 | 186.24 | 0.58 | 56.25 | 8.13 |
| *Neofusicoccum kwambonambiense*<br>CBS123642 | 1717 | 2871 | 44.21 | 44.06 | 210/61.88 | 352/34.6 | 387.07 | 262.94 | 0.58 | 56.33 | 7.62 |
| *Neofusicoccum parvum*<br>CBS123649 | 2185 | 6686 | 42.16 | 41.76 | 312/39.13 | 995/12.5 | 439.22 | 92.09 | 0.40 | 56.65 | 4.70 |
| *Neofusicoccum parvum* CMW9080 | 5188 | 5739 | 41.41 | 41.39 | 830/14.81 | 897/13.55 | 99.22 | 80.57 | 0.04 | 56.65 | 4.82 |
| *Neofusicoccum ribis* CBS115475 | 1994 | 3261 | 43.18 | 43.03 | 301/42.08 | 474/26.80 | 231.82 | 186.29 | 0.43 | 55.96 | 7.51 |
| *Neofusicoccum ribis* CBS121.26 | 2417 | 3145 | 43.12 | 43.03 | 245/51.65 | 371/33.92 | 280.49 | 167.59 | 0.51 | 56.03 | 7.35 |
| *Neofusicoccum umdonicola*<br>CBS123644 | 1343 | 2424 | 42.29 | 42.15 | 165/73.88 | 345/36.48 | 422.57 | 210.10 | 0.67 | 56.75 | 5.29 |

Twenty-six *Botryosphaeriaceae* genomes were annotated by predicting protein-coding genes with MAKER using BRAKER trained profiles. BUSCO analysis (Table 3) indicated that all *Botryosphaeriaceae* genomes had a high degree of completeness (average of 98.%, minimum of 95.1%). The genome annotations that were generated also had a high BUSCO completeness score (average of 97.8.%, minimum of 94.3%). When comparing theses BUSCO results to those of species with existing genome annotations on NCBI/JGI, it was clear that in five out of the six cases the genome annotations from the present study were more complete than others.

**Table 3.**
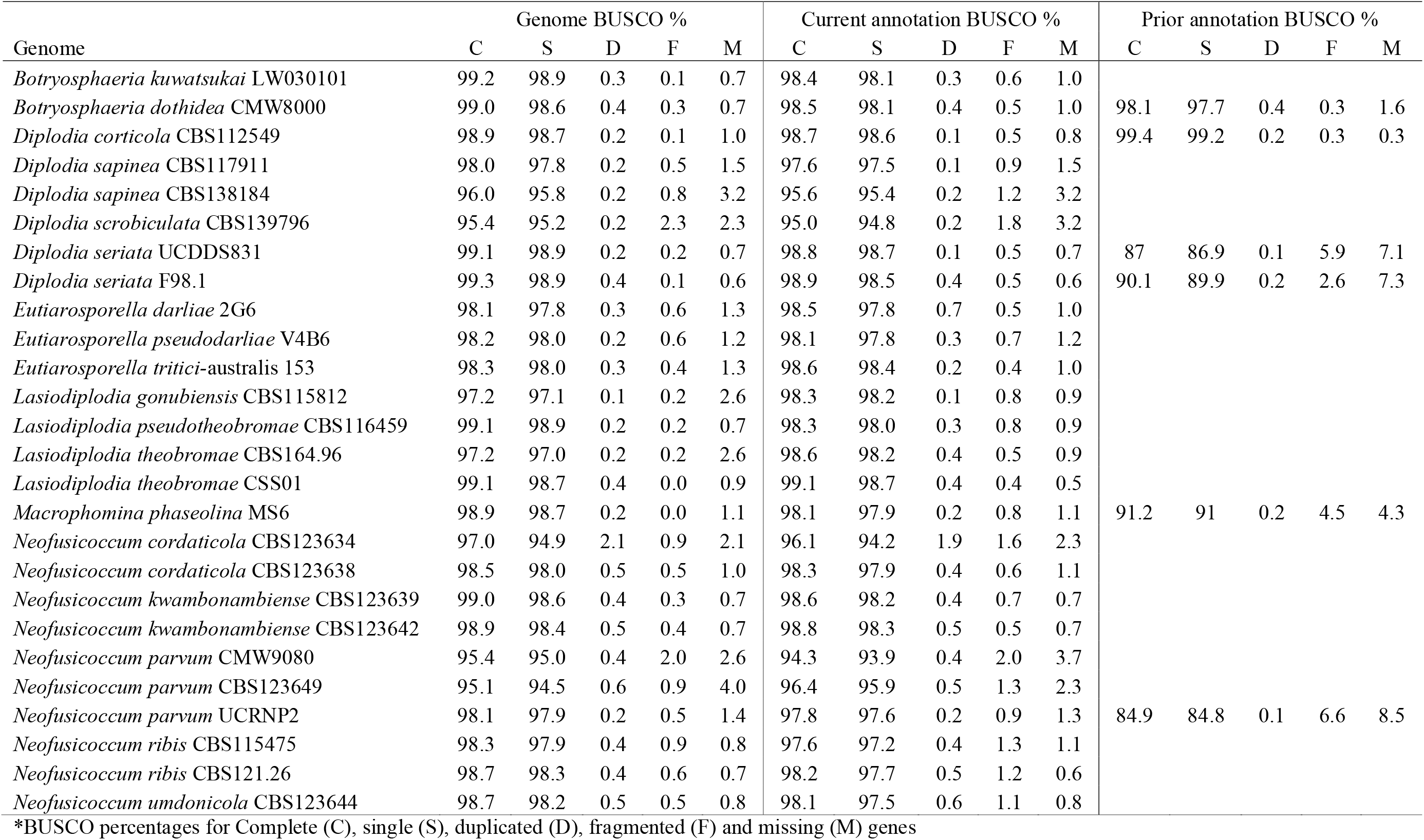
Genome and genome annotation completeness assesment*

### 3.2. Phylogenomic analyses

We identified 207 core orthologous genes from the collection of 26 *Botryosphaeriaceae*, 39 other Dothideomycetes and the outgroup (*Aspergillus nidulans*) genomes. Only orthologous genes that were represented by a single gene per species were retained. The results of the phylogenomic analyses corresponded well to previous phylogenies for the Dothideomycetes (Schoch *et al*. 2009; Schoch *et al*. 2006; Zhang *et al*. 2011). The phylogeny indicated the early divergence of the Dothideomycetidae (Dothideales, Capnodiales and Myrangiales) from the lineage containing the Pleosporomycetidae (Pleosporales, Hysteriales and Mytilinidiales) and other Dothideomycetes without current subclass designation (Figure 1). This phylogeny further supported the early divergence of the Botryosphaeriales from the ancestral Dothideomycetes lineage after the divergence of the Dothideomycetidae and Venturiales. The phylogenetic relationships between the *Botryosphaeriacaeae* were well defined and the phylogenetic placement of species and genera corresponded with that found in previous studies (Slippers *et al*. 2013; Yang *et al*. 2017).

**Figure 1.**
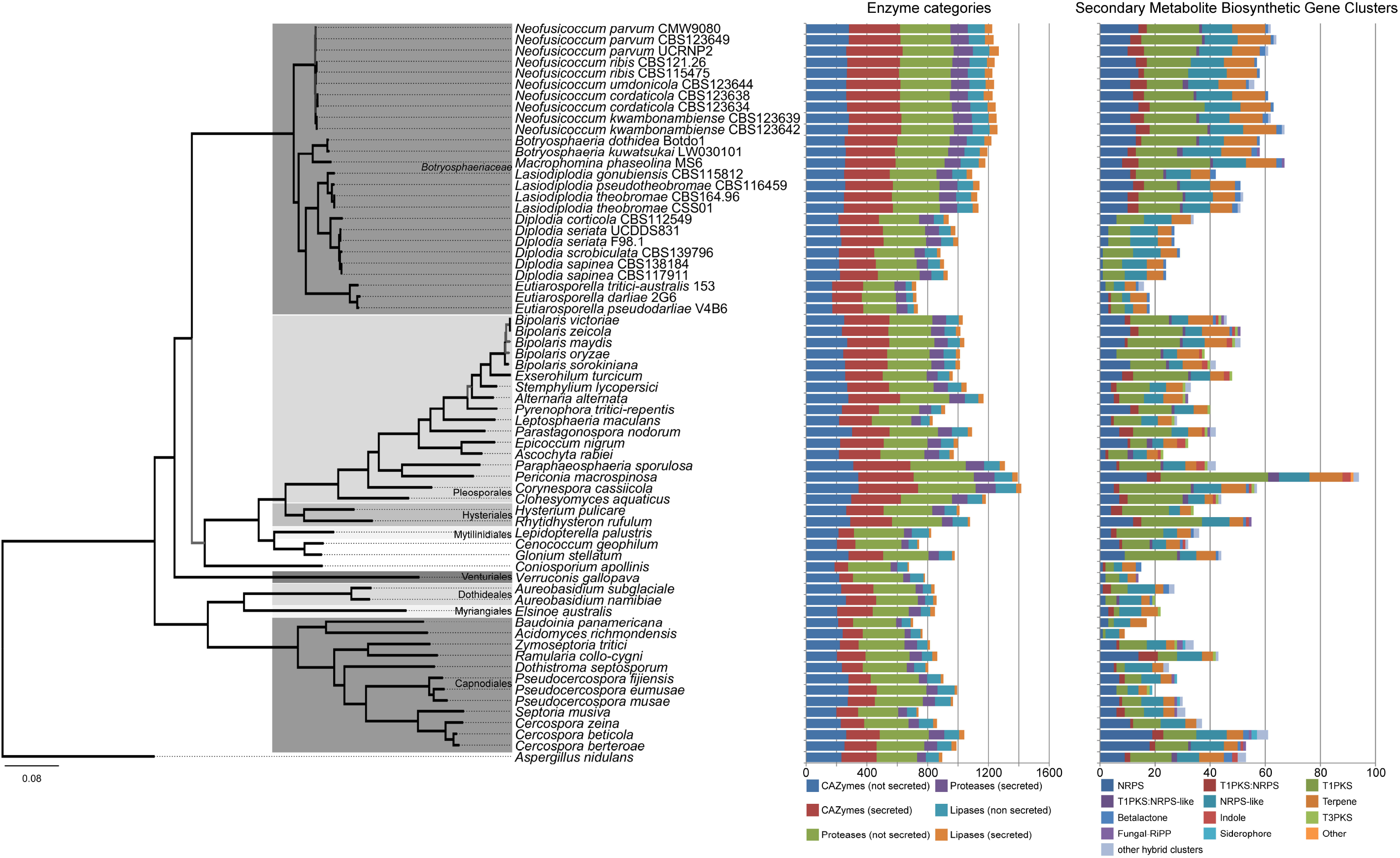
Phylogenomic tree and functional annotation summary of 26 *Botryosphaeriaceae*, 39 Dothideomycetes and one outgroup (*Aspergillus nidulans*). The supermatrix maximum likelihood phylogeny was determined using the sequence data of 207 single-copy core orthologous genes. Branches with 100% bootstrap support are indicated in black, those with less than 100% support are indicated in grey. The number of genes (total and secreted) annotated with CAZyme, proteases or lipase activity, as well as the number of gene cluster types involved in secondary metabolite biosynthesis are indicated using bar graphs.

### 3.3. Functional annotation

The genomes of the *Botryosphaeriaceae* differed noticeably in the number of predicted secreted proteins, hydrolytic enzymes and secondary metabolite BGCs they encode. However, distinct trends became apparent when considering the genus to which these genomes belong. Invariably, *Eutiarosporella* spp. had the fewest numbers of each of these functional annotation categories that we considered, followed by *Diplodia* spp. and *Botryosphaeria, Lasiodiplodia, Macrophomina, and Neofusicoccum* species had the most of these genes (Figure 1). This trend was also evident when considering the different classes of CAZymes/proteases/lipases and the different types of secondary metabolite BGCs (Figure 2).

**Figure 2.**
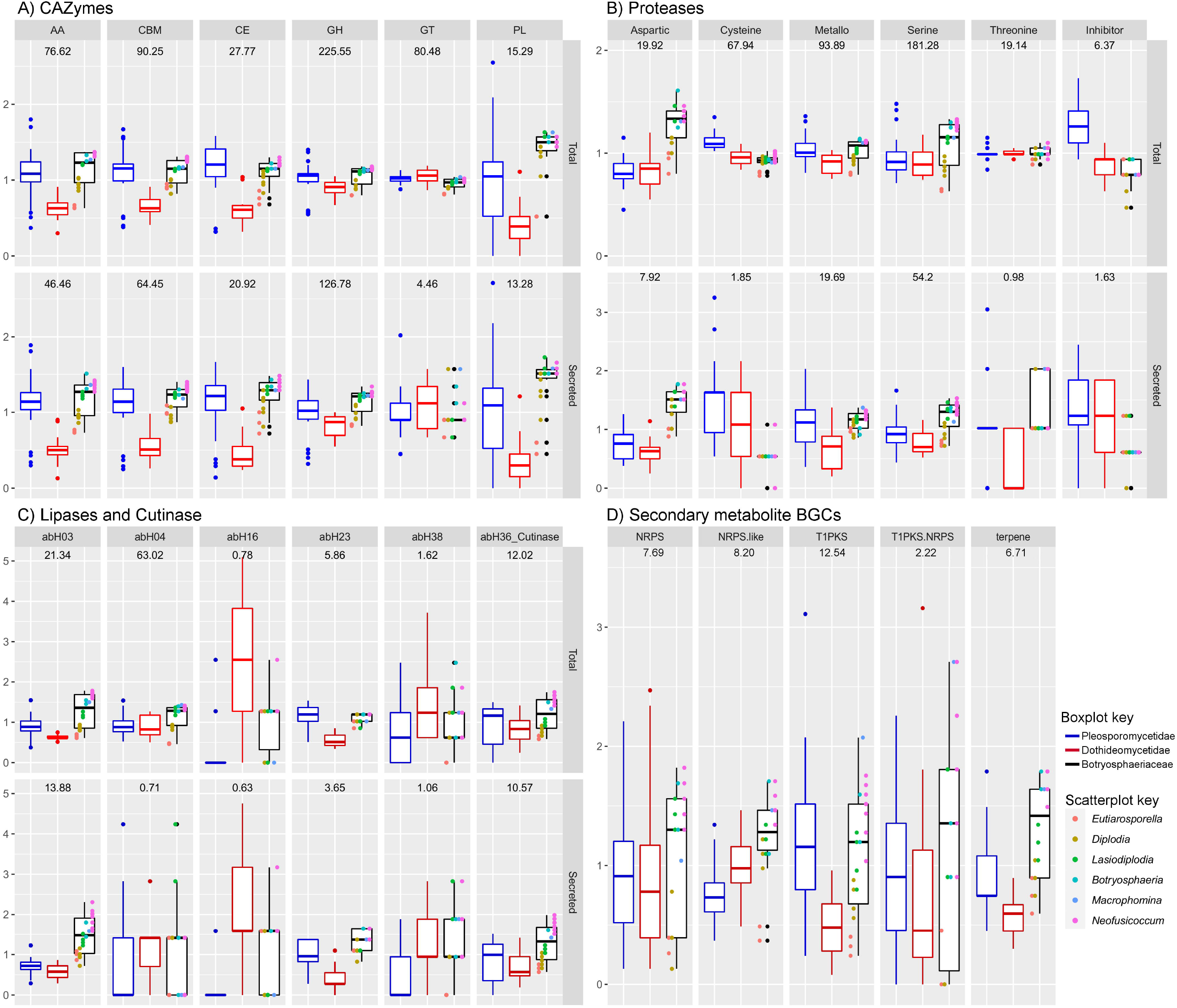
Box-and-whisker plots of the number of prominent CAZyme, protease and lipase families (total and secreted) and secondary metabolite BGC types present within the genomes of the considered *Botryosphaeriaceae* and other Dothideomycetes taxa. Taxa are placed into three categories: Dothideomycetidae, Pleosporomycetidae (plus related taxa without subclass designation) and *Botryosphaeriaceae*. The upper and lower bounds of the box represent the 1^st^ and 3^rd^ quartiles and the bar inside the median. The error lines (whiskers) represents 1.5 times the interquartile range (IQR) and outliers are indicated as dots. Additionally, the genera of the *Botryosphaeriaceae* are indicated by means of a scatterplot overlain on the *Botryosphaeriaceae* box-and-whisker plot.

The *Botryosphaeriaceae*, on average, had more of each CAZyme class than the Dothideomycetidae and a greater or roughly equal amounts than the Pleosporomycetidae, including the three *incertae sedis* taxa (Figure 2). The exception was for the glycosyltransferases (GT), where the *Botryosphaeriaceae* had fewer genes than the Dothideomycetidae and Pleosporomycetidae. The *Botryosphaeriaceae* were furthermore particularly rich in polysaccharide lyase (PL) genes, both the total number encoded in the genome and within the secretome. The most abundant secreted CAZyme families in the *Botryosphaeriaceae* were CBM1, AA3, GH3, GH43, GH5, AA9, CBM18, AA1, GH28 and CBM13 (Supplementary File 1).

The *Botryosphaeriaceae* had above-average numbers of aspartic-(A), metallo-(M) and serine-(S) proteases, especially in the species of *Botryosphaeria, Lasiodiplodia, Macrophomina, and Neofusicoccum* (Figure 2). These three protease classes were also the dominant proteases in the secretome. Furthermore the *Botryosphaeriaceae* had fewer than average number of cysteine (C) proteases and protease inhibitors (I) than the other Dothideomycetes. Notably, the *Botryosphaeriaceae* possessed a single secreted protease inhibitor family, namely I51.001 (serine carboxypeptidase Y inhibitor), whereas many other Dothideomycetes secreted protease inhibitors were of this family, as well as I09.002 (peptidase A inhibitor 1) or I09.003 (peptidase B inhibitor). *Diplodia sapinea* and *D. scrobiculata* had no secreted protease inhibitors. The most abundant secreted protease families among the *Botryosphaeriaceae* were S09, A01, S10, S08, M28, S53, S33, M43, M35, S12 (Supplementary File 1).

In the *Botryosphaeriaceae* and the Dothideomycetes, the most abundant lipases/lipase-like families were abH04 (Moraxella lipase 2 like), abH03 (Candida rugosa lipase-like), abH36, (cutinase) and abH23 (Filamentous fungi lipases) (Figure 2). The abH03, abH36 and abH23 lipase families were the main constituents of the predicted secretomes among the Dothideomycetes and the *Botryosphaeriaceae* had above-average numbers for these families.

The *Botryosphaeriaceae* genomes were rich in gene clusters involved in the synthesis of secondary metabolites. Type 1 polyketide synthases (t1PKS) were the most abundant type of gene cluster, followed by non-ribosomal peptide synthetases (NRPS) and NRPS-like, terpene synthases (TS) and t1PKS-NRPS hybrid clusters. The genomes of *Botryosphaeria, Lasiodiplodia, Macrophomina*, and *Neofusicoccum* contained higher than average numbers of t1PKS, NRPS-like, NRPS, and TS BGCs (Figure 3). Additionally, the *Botryosphaericeae* genomes contained betalactone BGCs (Supplementary File 1). The genomes of *Diplodia* and *Eutiarsporella* contained fewer than average of these gene clusters. Certain Dothideomycetes genomes, predominantly those of the Pleosporomycetidae also contained indole and type 3 PKS BGCs, however, these were not present in any of the *Botryosphaeriaceae* genomes.

**Figure 3.**
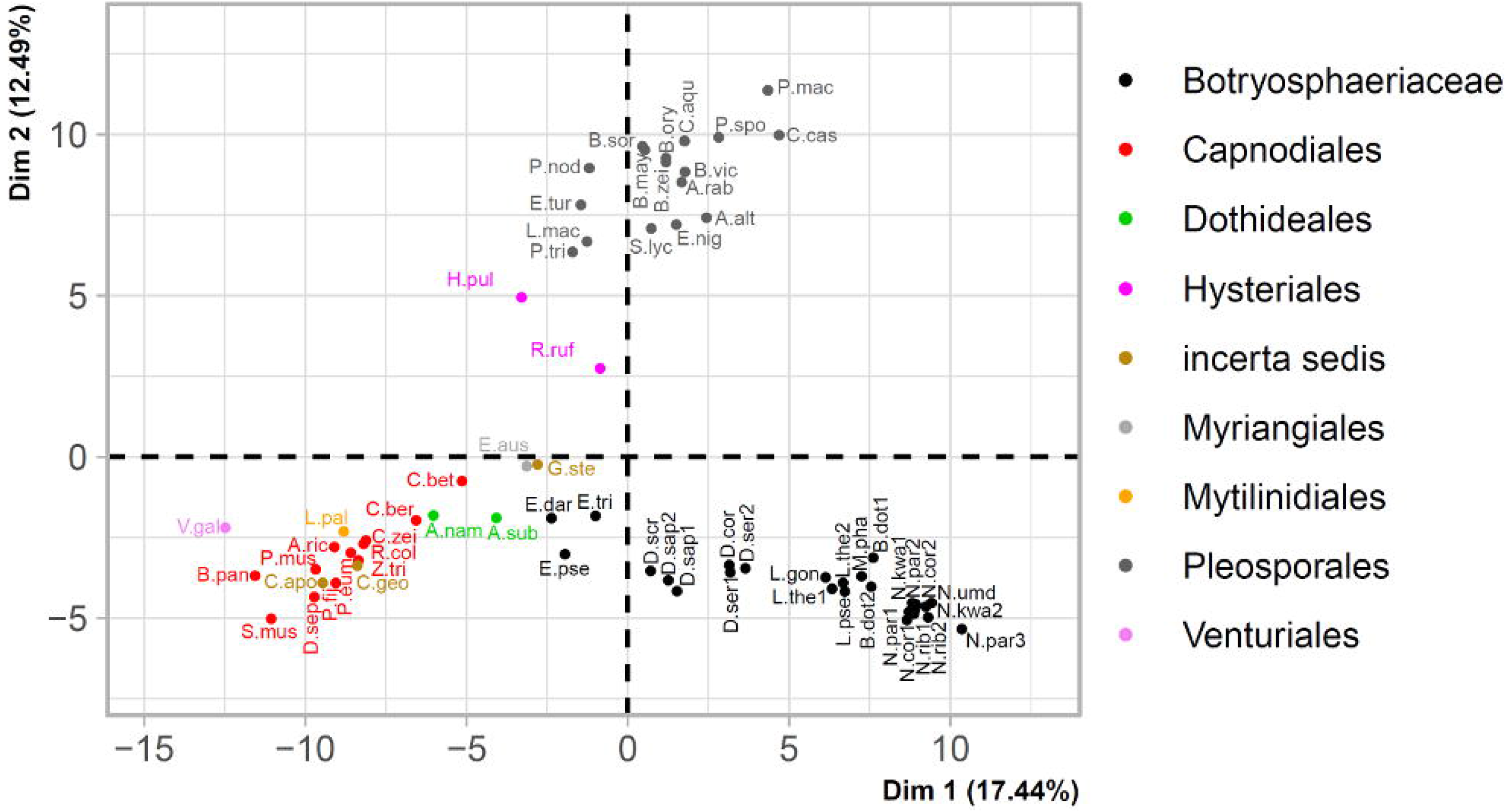
Principal component analysis of functional annotation categories of secreted proteins and secondary metabolite BGCs from *Botryosphaeriaceae* and other Dothideomycetes. Taxa are indicated using abbreviated names (Supplementary File 1) and colours indicate their Order/Family. The percentage variation accounted for by each principal component is indicated at each axis.

In the *Botryosphaeriaceae*, both genome size and the total gene number were strongly correlated with the number of secreted proteins, CAZymes, proteases, lipases and secondary metabolite gene clusters (Supplementary File 1). Within the Dothideomycetes, however, the numbers of secreted proteins, CAZymes, proteases, lipases, and secondary metabolite BGCs present within a genome were correlated to one another (i.e. species that contained large numbers of secreted proteins also contained large numbers of CAZymes, proteases, lipases and secondary metabolite BGCs), but only weakly correlated to genome size and the total number of genes (Supplementary File 1).

### 3.4. Gene family evolution

These results of the CAFE analyses (Supplementary file 2) indicated that expansions and contractions of CAZyme gene families occurred at roughly similar levels. This was after the divergence of the Botryosphaeriales ancestor from the Pleosporomycetidae until the formation of the *Botryosphaeriaceae* crown group (61 MYA). During this time, protease gene families apparently experienced more contractions than expansions, lipase gene families had slightly more expansions than contractions and secondary metabolite BGCs experienced a large amount of gene family contractions. Several CAZyme gene families (AA1, AA3, AA7, AA8, AA9, CBM1, CBM18, CE4, CE5, GH3, GH10, GH28, GH43, GH78, GT1, GT2, GT25, PL1 and PL3) experienced rapid expansion (i.e. greater than expected under the birth/death model of gene family evolution) prior to the divergence of the *Botryosphaeriaceae* crown group.

After the divergence of the *Botryosphaeriaceae*, the genera *Botryosphaeria, Lasiodiplodia, Macrophomina, and Neofusicoccum* experienced more gene family expansions than contractions, whereas the opposite was observed for the *Diplodia* and *Eutiariosporella*. Among the *Neofusicoccum* spp., the AA3, AA7, GH3 and GT2 gene families were rapidly expanding. Similarly, among the *Lasiodiplodia* spp., the AA7 and GH106 gene families were rapidly expanding. Conversely, among the *Diplodia* spp. the AA7 gene family was rapidly contracting, as were the AA1, AA3, AA7, GH28, PL1 and PL3 gene families among the *Eutiarosporella* spp.

### 3.5. Principal component analysis and hierarchical clustering

Hierarchical clustering separated the taxa into four groups (Supplementary File 3). The taxa in the Pleosporomycetidae and Dothideomycetidae generally clustered separately, however there was no overall clustering based on taxonomic placement. *Botryosphaeriaceae* species were present in three of the four dominant clusters.

A first cluster included *B. dothidea, B. kuwatsukai, M. phaseolina, L. theobromae, L. pseudotheobromae*, and *Neofusicoccum* spp., as well as several Pleosporales (*Alternaria alternata, Clohesyomyces aquaticus, Corynespora cassiicola Paraphaeosphaeria sporulosa* and *Periconia macrospinosa*). A second cluster mostly contained taxa from the Pleosporales (*Aschochyta rabiei, Bipolaris* spp., *Epicoccum nigrum* and *Stemphylium lycopersici*), but also contained *D. seriata, D. corticola* and *L. gonubiensis*. A third cluster contained taxa from both the Dothideomycetidae and Pleosporomycetidae, as well as *D. sapinea, D. scrobiculata* and *Eutiarosporella* spp. A fourth cluster was dominated by taxa from the Dothideomycetidae with the exception of *L. palustris, C. geophilum, C. apollinis* and *V. gallopava*.

PCA of the functional annotation categories clustered the data along 65 dimensions/principal components. The first two dimensions (Figure 3) accounted for 29.9% of the variance among the taxa. The first dimension accounted for 17.4% and the second dimension for 12.5% of the variance. The first dimension was most strongly influenced by a number of secreted CAZyme (AA3, CBM1, GH131, PL3, CBM13, CBM18, CE8, CE12, AA7, GH43, PL4, PL1, CE5 and CBM63), cutinase (abH36) and lipase (abH03 and abH23) and protease (S09 and A01) families, as well as terpene BGCs. The second dimension was most strongly influenced by CAZyme (GH145, CBM3, GH6, GH11, PL26, CBM60, GH7, AA12, AA9, CBM2, GH16, CBM6, CBM87 and CE18), protease (S01, M14 and M36) families, as well as the indole and T3PKS BGC types.

The *Botryosphaeriaceae* were distributed mainly along the first dimension of the PCA and clustered into three groups. *Eutiarosporella* spp. clustered at the lower ranges of the first dimension (x-axis) followed by clusters accomodating the *Diplodia* species towards the middle ranges and the other genera of *Botryosphaeriaceae* clustered at the high ranges of the x-axis. The *Botryosphaeriaceae* clustered along a relatively narrow range along the second dimension (y-axis) compared to the other Dothideomycetes. The clustering of the other Dothideomycetes along the x-axis was correlated to their clustering on the y-axis: taxa towards the higher end of the x-axis also occurred towards the higher end of the y-axis. The clustering of taxa did not correspond to their nutritional lifestyle, but their taxonomic placement was reflected in their clustering.

### 3.6. Genome Architecture

Two-dimensional heatmaps of the 5’ and 3’ FIRs of the 26 *Botryosphaeriaceae* genomes indicated no genome compartmentalization (Figure 4 and Supplementary File 4). This was evident from the unimodal gene density distributions of these genomes. Genomes of *Botryosphaeria, Lasiodiplodia and Macrophomina* had a higher proportion of genes in gene sparse regions than the other species in this family. *Eutiarosporella* spp., *D. sapinea, D. scrobiculata* and *Neofusicoccum* spp. had fewer genes in gene sparse regions.

**Figure 4.**
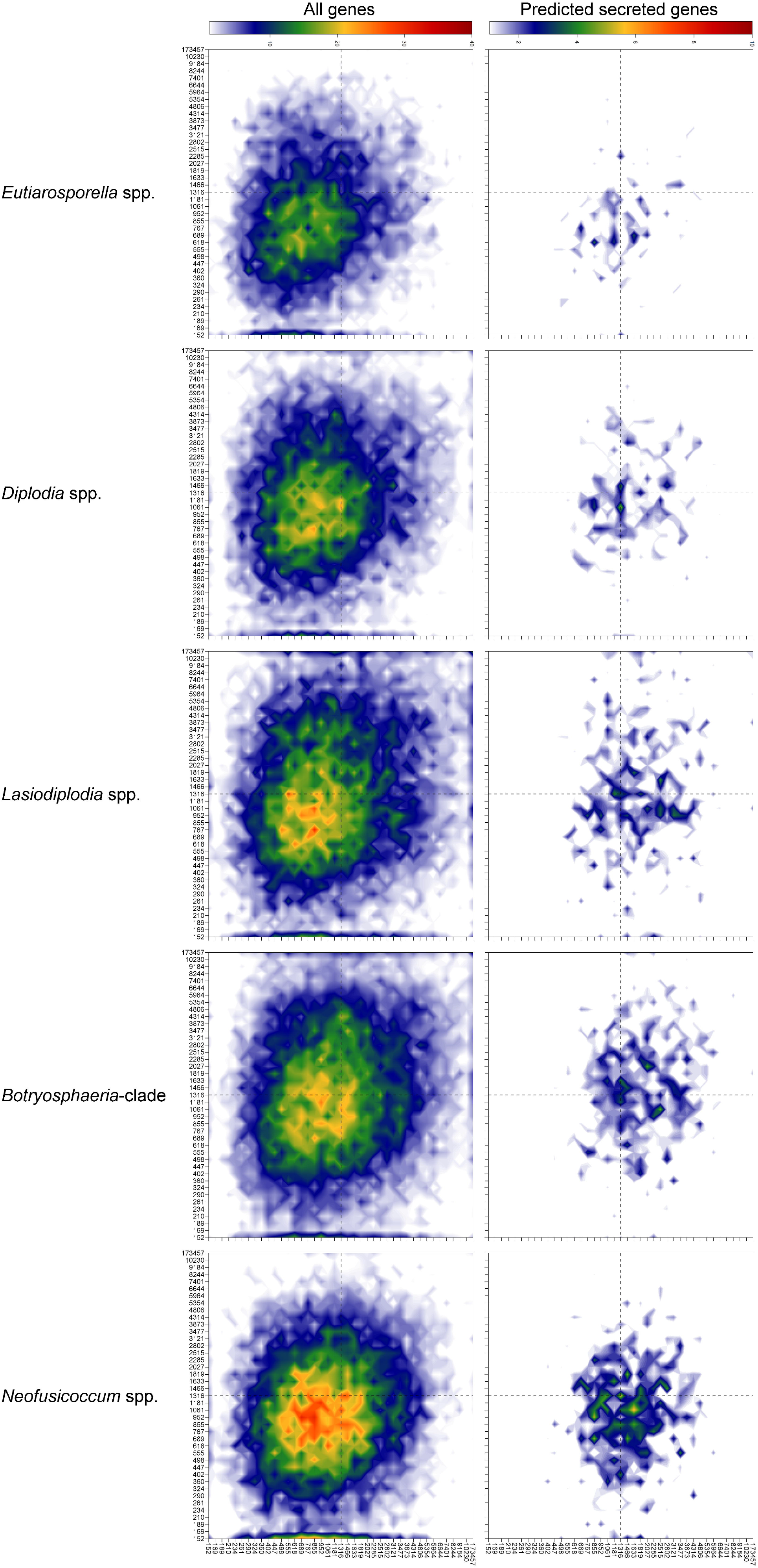
Gene density landscape of the genera of *Botryosphaeriaceae*. The two-dimensional heatmap shows the distribution of genes to gene dense or sparse regions of the genome based on their 5’ and 3’ flanking intergenic regions (FIRs). Two-dimensional heatmaps of each genus is the average across bins of all genomes in the group. The heatmaps labelled as “*Botryosphaeria-*clade” include the genomes of *B. dothidea, B. kuwatsukai* and *M. phaseolina*. The values on the axes are distances in base pairs and signify the upper limit of each bin. The colours of the heatmap represents the number of genes present within each two-dimensional bin.

The predicted secreted proteins of the *Botryosphaeriaceae* contained a greater number of genes in gene sparse regions than the total predicted genes (Supplementary File 4). The total CAZymes contained a higher proportion of genes in gene sparse regions than the total predicted genes. The secreted CAZyme gene density distribution was very similar to that of the total CAZymes. The secreted lipases and cutinases, however, occurred more frequently in gene sparse regions than the total lipases and cutinases (Supplementary File 4). This trend was also observed for the secreted proteases, although not as strongly. Genes associated with secondary metabolite BGCs were less prevalent in gene sparse regions.

The levels of repetitive sequences for most *Botryosphaeriaceae* genomes were between 3 and 8 percent of the total genome size (Table 4). The two *Botryosphaeria* spp. differed considerably in their repeat content (3.48 vs. 11.88%). The genome of *M. phaseolina* also had a higher than average repeat content (16.37%). Among the *Neofusicoccum* species the genomes of *N. parvum* and *N. umdonicola* had less repetitive sequences than the other genomes of this genus. The genomes of *D. sapinea* and *D. scrobiculata* also contained less repetitive sequences than the other two *Diplodia* species.

**Table 4.** Summary of genomic architecture features of *Botryosphaeriaceae* genomes

|  | Genome size (Mb) | Repetitive sequences (bp) | % of repetitive sequences | Genome GC % | % genome that is GC rich (>50%) | Genes in TA rich regions | Secreted genes in TA rich regions | % genome affected by RIP | Number of LRARs | Total size (bp) of all LRARs |
| --- | --- | --- | --- | --- | --- | --- | --- | --- | --- | --- |
| *Eutiarospora darliae* 2G6 | 27.27 | 1 415 283 | 5.19 | 61 | 98.3 | 3 | 0 | 0.91 | 4 | 32 276 |
| *E. pseudodarliae* V4B6 | 26.74 | 1 290 421 | 4.83 | 61.53 | 98.6 | 7 | 0 | 0.4 | 0 | 0 |
| *E. tritici-australis* 153 | 26.59 | 1 544 103 | 5.81 | 61.66 | 97.3 | 0 | 0 | 2.54 | 14 | 95 508 |
| *Diplodia sapinea* CBS117911 | 36.05 | 1 344 948 | 3.73 | 56.84 | 95 | 73 | 4 | 1.12 | 2 | 9 500 |
| *D. sapinea* CBS138184 | 35.24 | 1 305 464 | 3.7 | 56.73 | 94.7 | 68 | 6 | 0.96 | 2 | 12 500 |
| *D. scrobiculata* CBS139796 | 34.93 | 1 110 716 | 3.18 | 57.01 | 95.6 | 63 | 6 | 0.82 | 1 | 13 000 |
| *D. seriata* UCDDS831 | 37.27 | 1 711 911 | 4.61 | 56.6 | 96.8 | 29 | 2 | 2.63 | 21 | 125 537 |
| *D. seriata* F98.1 | 37.12 | 1 593 160 | 4.27 | 48.47 | 94.5 | 51 | 1 | 1.87 | 34 | 554 000 |
| *D. corticola* CBS112549 | 34.99 | 2 102 085 | 6.01 | 47.92 | 94.5 | 31 | 3 | 3.39 | 42 | 690 014 |
| *Lasiodiplodia theobromae* CBS164.96 | 42.97 | 1 498 597 | 3.49 | 54.74 | 89.9 | 354 | 32 | 0.72 | 3 | 59 500 |
| *L. theobromae* CSS01 | 43.28 | 1 543 231 | 3.57 | 48.15 | 89.8 | 306 | 33 | 1.04 | 18 | 375 000 |
| *L. pseudotheobromae* CBS116459 | 43.01 | 1 402 977 | 3.26 | 54.66 | 89.2 | 377 | 36 | 0.92 | 12 | 375 000 |
| *L. gonubiensis* CBS115812 | 41.14 | 1 330 240 | 3.23 | 54.72 | 92.1 | 223 | 24 | 1.18 | 12 | 345 500 |
| *Botryosphaeria dothidea* CMW8000 | 43.5 | 1 515 197 | 3.48 | 54.3 | 89.8 | 385 | 53 | 0.88 | 1 | 5 000 |
| *B. kuwatsukai* LW030101 | 47.39 | 5 628 118 | 11.88 | 53.09 | 81.1 | 406 | 41 | 9.65 | 170 | 1 387 657 |
| *Macrophomina phaseolina* MS6 | 48.88 | 8 000 688 | 16.37 | 52.33 | 80.2 | 228 | 23 | 12.92 | 172 | 2 658 966 |
| *Neofusicoccum cordaticola* CBS123634 | 45.71 | 3 594 350 | 7.86 | 54.9 | 86.5 | 566 | 54 | 3.7 | 105 | 1 071 347 |
| *N. cordaticola* CBS123638 | 43.56 | 3 614 552 | 8.3 | 55.92 | 88.7 | 393 | 40 | 3.99 | 47 | 324 966 |
| *N. kwambonambiense* CBS123639 | 44.17 | 3 369 216 | 7.62 | 55.92 | 89.1 | 408 | 40 | 2.75 | 64 | 693 395 |
| *N. kwambonambiense* CBS123642 | 44.21 | 3 592 960 | 8.13 | 56.04 | 89.2 | 459 | 49 | 2.35 | 56 | 554 628 |
| *N. parvum* CMW9080 | 41.41 | 1 944 617 | 4.7 | 56.54 | 91.3 | 406 | 40 | 1.36 | 11 | 63 557 |
| *N. parvum* CBS123649 | 42.16 | 2 032 314 | 4.82 | 56.06 | 91.8 | 360 | 36 | 1.65 | 14 | 88 222 |
| *N. parvum* UCRNP2 | 42.52 | 2 167 908 | 5.1 | 56.76 | 91.4 | 407 | 35 | 1.59 | 11 | 72 874 |
| *N. ribis* CBS115475 | 43.18 | 3 241 261 | 7.51 | 55.71 | 89.6 | 379 | 38 | 4.52 | 79 | 911 957 |
| *N. ribis* CBS121.26 | 43.12 | 3 171 232 | 7.35 | 55.88 | 89.1 | 423 | 51 | 4.71 | 25 | 153 638 |
| *N. umdonicola* CBS123644 | 42.29 | 2 237 372 | 5.29 | 56.51 | 91.6 | 415 | 48 | 1.38 | 11 | 76 232 |

On average, the *Botryosphaeriaceae* genomes had less than 10% of their genomes composed of TA rich regions (GC<50%) (Table 4). The genomes of *B. kuwatsukai* LW030101 and *M. phaseolina* had 18.9% and 19.8% of their genomes as TA rich regions, respectively. The genomes of *Diplodia* and *Eutiarosporella* had fewer TA rich regions (approximately 5% of the genome) compared to the other genera. Less than 5% of genes were present in TA rich regions and secreted genes were not found to be over-represented among these genes (Supplementary file 4). The genomes of *Diplodia* and *Eutiarosporella* had considerably fewer genes associated with TA rich regions, than the other taxa of this family.

Analysis of the prevalence of RIP in the genomes of *Botryosphaeriaceae* indicated that this has occurred to varying degrees in these genomes (Table 4). *Neofusicoccum* spp. had between 1.36 and 4.71 % of their genome affected by RIP. The level of RIP was similar between different genomes of the same *Neofusicoccum* species. *Neofusicoccum parvum* and *N. umdonicola* had lower proportions of RIP affected sequences than the other species of the genus. There was a large (>10-fold) difference in the level of RIP between *B. dothidea* and *B. kuwatsukai*. The genome of *M. phaseolina* had the highest (12.92 %) amount of RIP of all the *Botryosphaeriaceae. Lasiodiplodia* spp. had RIP levels between 0.72 and 1.18 %. The level of RIP in the genomes of *D. sapinea* and *D. scrobiculata* was lower than in *D. seriata* and *D. corticola*. The genome of *E. tritici-australis* had more than double the level of RIP affected regions than the other two species of the genus.

## 4. Discussion

This study represents the first large-scale comparative genomics-level consideration of all available genomes of *Botryosphaeriaceae*. The results showed that the included *Botryosphaeriaceae* genomes, especially those of *Botryosphaeria, Macrophomina, Lasiodiplodia*, and *Neofusicoccum*, encode high numbers of secreted hydrolytic enzymes and secondary metabolite BGCs. This apparently emerges due to these fungi having large numbers of genes belonging to enzyme classes and secondary metabolite types involved in interactions with plants. The results also indicate that the *Botryosphaeriaceae* are most similar to species of the Pleosporamycetidae based on secreted enzyme and secondary metabolite profiles and that nutritional lifestyle could not be deduced from the data. *Botryosphaeriaceae* genomes were furthermore determined not be be compartmentalized based on gene density or GC-content.

There was a strong correlation between the number of hydrolytic enzymes and secondary metabolite BGCs, and the genome size and gene number of the *Botryosphaeriaceae* considered in this study. This correlation between genome size and gene number has generally not been seen in other fungi (Mohanta and Bae 2015; Ohm *et al*. 2012; Raffaele and Kamoun 2012) because transposable elements and repetitive DNA vary significantly among species (Raffaele and Kamoun 2012). A recent comparison of Dothideomycetes genomes (Haridas *et al*. 2020) also showed that genome size and gene number were not correlated to the abundance of functional annotation classes; neither to the lifestyle or phylogenetic placement of a species.

The genomes of *Botryosphaeria, Macrophomina, Lasiodiplodia*, and *Neofusicoccum* had abundant secreted hydrolytic enzyme and secondary metabolite BGCs. This is a pattern that is most similar to prominent necrotrophic plant pathogens (*A. alternata, C. casiicola*), saprobes (*C. aquaticus, P. sporulosa*) and the endophyte/latent pathogen *P. macrospinosa* in the Pleosporales. The pattern consistent with reports that necrotrophic pathogens tend to have higher numbers of hydrolytic enzymes and secondary metabolite toxins than biotrophs and symbiotic fungi (Lyu *et al*. 2015; Zhao *et al*. 2014). An abundance of secreted hydrolytic enzymes and secondary metabolite BGCs found in the *Botryosphaeriaceae* is also similar to that of other species of woody endophytes. Studies on such endophytic species have shown that they have similar or higher amounts of various secreted enzymes (notably plant cell wall degrading enzymes) or secondary metabolites than closely related plant pathogenic species (Hacquard *et al*. 2016; Schlegel *et al*. 2016; Xu *et al*. 2014; Yang *et al*. 2019). The specific gene families that are enriched, however, differ among endophytic lineages, due to the evolutionary independent origins of endophytism (Hacquard *et al*. 2016; Schlegel *et al*. 2016; Xu *et al*. 2014; Yang *et al*. 2019). It has furthermore been noted that fungi with dual lifestyles (e.g. fungi with endophytic and pathogenic phase) have large numbers of CAZymes (Queiroz and Santana 2020; Wang *et al*. 2015). These observations also emerging from the present study are consistent with the known lifestyle of *Botryosphaeriaceae* as latent pathogens.

The *Botryosphaeriaceae* genomes were rich in CAZymes, especially those involved in plant cell wall degradation (PCWD), although at lower numbers in the genomes of *Diplodia* and *Eutiarosporella* species. CAZymes involved in the degradation of cellulose, hemicellulose and pectin were present in all *Botryosphaeriaceae*. The genomes of *Botryosphaeria, Macrophomina, Lasiodiplodia* and *Neofusicoccum* were particularly rich in CAZyme families involved in cell wall degradation. Specifically, CAZymes involved in plant, general and fungal cell wall degradation (Levasseur *et al*. 2013; Ohm *et al*. 2012; Zhao *et al*. 2013) were abundant in the genomes of the above-mentioned genera.

The *Botryosphaeriaceae* secretomes were rich in CAZyme families involved in the recognition of cellulose (CBM1) and chitin (CBM18). Although carbohydrate-binding domains have no catalytic activity of their own they play important roles in substrate recognition and binding of other CAZymes (Boraston *et al*. 2004; Christiansen *et al*. 2009), they are also involved in the protection of fungal cell walls from degradation by host enzymes and prevention of host detection (De Jonge and Thomma 2009; Marshall *et al*. 2011). High numbers of secreted CAZymes involved with PCWD have also been found in previous studies of *N. parvum* and *D. seriata* (Massonnet *et al*. 2016; Morales-Cruz *et al*. 2015) and in other, especially necrotrophic, Dothideomycetes (Lopez *et al*. 2018; Manning *et al*. 2013; Verma *et al*. 2016; Zeiner *et al*. 2016; Zhao *et al*. 2013). The abundance of these CAZyme families in some genera of *Botryosphaeriaceae* suggests that cell wall degradation plays an important role in the biology of these fungi.

Several important CAZyme families that are common among Dothideomycetes were absent from all the *Botryosphaeriaceae* genomes, i.e. Acetyl xylan esterase (CE3) (Biely *et al*. 1985), Pyrroloquinoline quinone-dependent oxidoreductase (AA12) (Takeda *et al*. 2015), endo-α-1,4-polygalactosaminidase (GH114) (Bamford *et al*. 2019) and α-L-arabinofuranosidase/β-xylosidase (GH54) (Miyanaga *et al*. 2006). The absence of these CAZyme families in the *Botryosphaeriaceae*, does not necessarily indicate a gap in the metabolic repertoire of these fungi because a large degree of functional redundancy is commonly seen in fungal CAZyme repertoires (Couger *et al*. 2015; Couturier *et al*. 2016; Goulet and Saville 2017). Interestingly, some of the CAZyme families that can functionally compensate for the absence of the above-mentioned CAZyme families are those that were found to be among the most abundant secreted CAZyme families of the *Botryosphaeriaceae* (e.g. CE16, AA3, AA7, GH15 and GH3).

The *Botryosphaeriaceae* were rich in secreted serine-, metallo- and aspartic-proteases. Protease families (A01, S08, S09, S10) that were previously identified as the most common secreted proteases among Dothideomycetes (Ohm *et al*. 2012) were also the most abundant in the *Botryosphaeriaceae*. Secreted proteases play important roles in nutrient acquisition, signalling and degradation of plant defences (Carlile *et al*. 2000; Olivieri *et al*. 2002; Plummer *et al*. 2004; Thon *et al*. 2002). Although secreted proteases are abundant in several necrotrophic pathogens, e.g. *Corynespora cassiicola* (Lopez *et al*. 2018) and several *Colletotrichum* spp. (Baroncelli et al. 2016), no patterns between nutritional lifestyle and the abundance of secreted proteases could be distinguished. The precise function of most of these proteases in the *Botryosphaeriaceae* are unknown and their role during infection and disease expression remains to be determined.

A lower abundance and diversity of secreted protease inhibitors of the *Botryosphaeriaceae* suggests a reduced capacity and/or need for extracellular enzyme inhibition. Plant pathogenic fungi secrete protease inhibitors to inhibit plant proteases involved in defence responses (Jashni *et al*. 2015) and several protease inhibitors are known virulence factors, e.g. *avr2* of *Cladosporium fulvum* (Rooney *et al*. 2005; van Esse *et al*. 2008) and *Pit2* of *Ustilago maydis* (Mueller *et al*. 2013). However, the exact role of many fungal protease inhibitors, such as those secreted by the *Botryosphaeriaceae* and Dothideomycetes, remains unknown (Dunaevsky *et al*. 2014).

*Botryosphaeriaceae*, especially species of *Botryosphaeria, Macrophomina, Lasiodiplodia* and *Neofusicoccum* possessed high numbers of secreted lipases. Three lipase families were present in high numbers in the secretomes of the Dothideomycetes, i.e *Candida rugosa* lipase-like (abH03), cutinases (abH36 and CE5) and Filamentous fungi lipases (abH23). The *Botryosphaericeae* genomes were rich in secreted enzymes for these three families. Lipases and cutinases are important for fungal penetration of host tissue (Kolattukudy 1985; Voigt *et al*. 2005), growth and adhesion (Feng *et al*. 2009; Gácser *et al*. 2007) and manipulation of host defences (Christensen and Kolomiets 2011). The abundance of these secreted lipases and cutinases emphasises the importance of their role during the infection process in the *Botryosphaeriaceae*.

The genomes of *Botryosphaeria, Macrophomina, Lasiodiplodia* and *Neofusicoccum* contained many BGCs, especially t1PKS, NRPS, NRPS-like and TS type clusters. The products produced by most of these clusters are unknown, however, the products of some clusters could be determined. These compounds included melanin, phytotoxins (ACT-Toxin II, (-)-Mellein), siderophores (dimethylcoprogen) and antioxidants (pyranonigrin E). The ability to produce secondary metabolite toxins have been associated with pathogenic fungi’s lifestyle, host range and virulence (Ito *et al*. 2004; Ohm *et al*. 2012; Stergiopoulos *et al*. 2013). Many plant pathogenic fungi have large numbers of secondary metabolite BGCs, e.g. *Bipolaris* spp. (Zaccaron and Bluhm 2017), *Corynespora cassiicola* (Lopez *et al*. 2018), *Colletotrichum* spp (Baroncelli et al. 2016) and *Pyrenophora teres* (Ohm *et al*. 2012), but so also do fungi with other lifestyles such as the saprobic *Annulohypoxylon stygium* (Wingfield *et al*. 2018), *Hysterium pulicare*, and *Rhytidhysteron rufulum* (Ohm *et al*. 2012). Despite the observation that the total abundance of secondary metabolite BGCs does not predict lifestyle, several fungal toxins are able to modulate a fungal species’ host range or virulence, e.g. the AF-toxin of *A. alternata* (Ito et al. 2004), the *Hybrid-1,2* and *3* genes of *Eutiarosporella darliae* and *E. pseudodarliae* (Thynne et al. 2019) and the HC-toxins of *Bipolaris zeicola* (Wolpert et al. 2002).

The *Botryosphaeriaceae* genera were shown to possess non-compartmentalized genomes. These species were characterized as having moderate to high %GC, RIP-affected genomes with low amounts of repetitive DNA, a slight preferential localization of secreted genes to gene sparse regions and no preferential localization of secreted genes to TA rich regions. Most Dothideomycetes do not have compartmentalized genomes, but several important pathogenic species (e.g. *L. maculans* and *Pseudocercospora* spp.), have genomes with high levels of repetitive DNA (often transposable elements) (Chang *et al*. 2016; Rouxel *et al*. 2011) enriched for fast-evolving genes related to pathogenicity or virulence (Chang *et al*. 2016; Grandaubert *et al*. 2014; Testa *et al*. 2016). However, not all rapidly evolving fungal phytopathogens have this characteristic ‘two-speed’ genome architecture (Frantzeskakis *et al*. 2019). Where ‘two-speed’ genomes rely on the action of leaky RIP to generate variation for selection to act on, ‘one-speed’ genomes of fast-evolving plant pathogens rely on the absence of RIP that allows gene duplication/copy number variation to generate variation (Frantzeskakis *et al*. 2019). The *Botryosphaeriaceae* are not like those species with ‘two-speed’ genomes as they don’t have compartmentalized genomes, but RIP is also not completely absent as seen in species with ‘one-speed’ fast-evolving genomes.

This study is the first large-scale comparative genomics study to consider all available genomes of *Botryosphaeriaceae*. It has illustrated large variability in the secreted hydrolytic enzyme and secondary metabolite biosynthetic repertoire between genera of this family. Furthermore, we have demonstrated similarities between the *Botryosphaeriaceae* and necrotrophic plant pathogens and endophytes of woody plants, emphasising their role as latent pathogens. This study highlights the importance of these genes in the infection biology of *Botryosphaeriaceae* species and their interaction with plant hosts. This knowledge will be useful in future studies aimed at understanding the mechanisms of endophytic infections and how these transition to a pathogenic state. The results should also help to better understand the genetic factors involved in determining the complex question of host range in the Botryosphaerianceae.

## 5. Acknowledgements

We are grateful to the University of Pretoria, The Department of Science and Technology (DST)/National Research Foundation (NRF) Centre of Excellence in Tree Health Biotechnology and members of the Tree Protection Cooperative Program for financial support of this study. The authors acknowledge the Centre for Bioinformatics and Computational Biology, University of Pretoria and the Centre for High Performance Computing (CHPC), South Africa for providing computational resources to this research project. Lastly, our sincere thanks to Alisa Postma for her guidance during the genome assembly and annotation process and to Dr Tuan Duong for his critical comments on this manuscript.

---

**Supplementary File 1:** Excel file of the functional annotation data

**Supplementary File 2:** Phylogenetic tree indicating the gene family contractions/expansions of CAZymes, proteases, lipases and secondary metabolite BGCs among the *Botryosphaeriaceae* and other Dothideomycetes

**Supplementary File 3:** Hierarchical clustering and heatmap of *Botryosphaeriaceae* and other representative Dothideomycetes species based on the number of functional annotation categories of secreted and secondary metabolite BGCs. Overrepresented (red and dark red) and underrepresented (blue and dark blue) values are scaled relative to the column mean.

**Supplementary File 4:** Excel file containing data of gene association with gene-sparse/TA-rich regions

