## Supplementary figures and images for "Abundant secreted hydrolytic enzymes and secondary metabolite gene clusters in genomes of the *Botryosphaeriaceae* reflect their role as important plant pathogens"

### Supplementary File 2

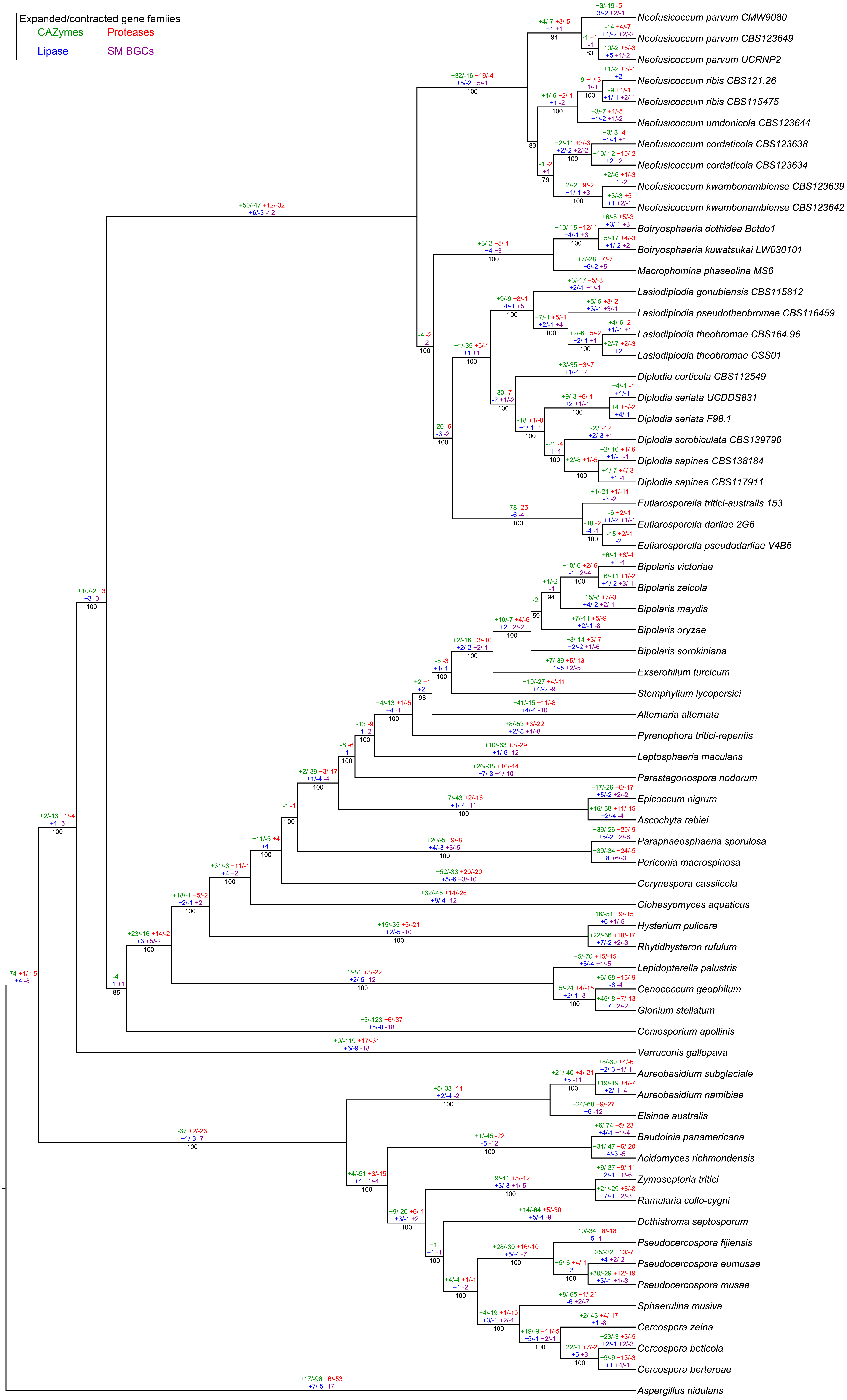

### Supplementary File 3

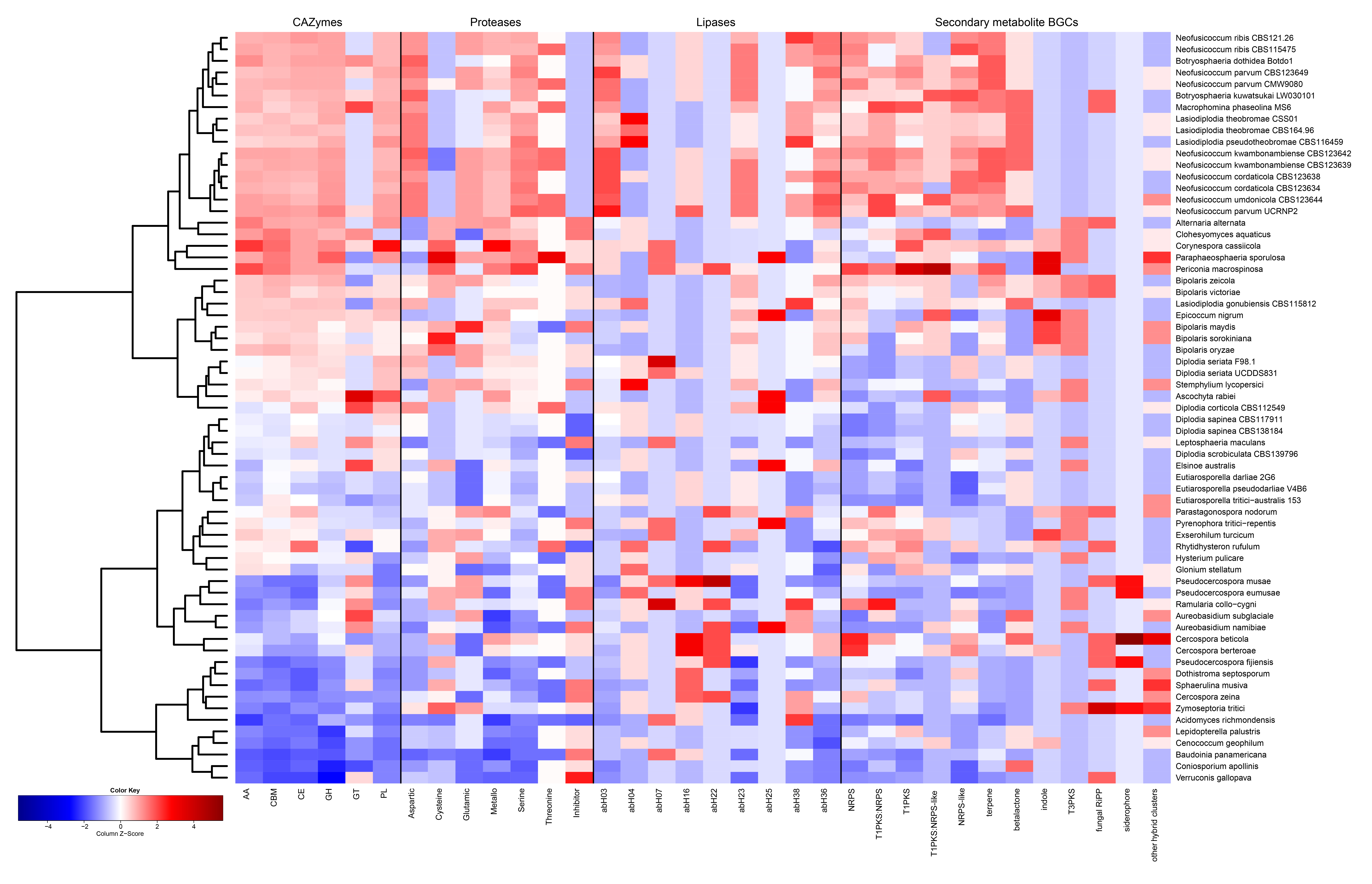
